# Daphnids Can Safeguard the Use of Alternative Bioassays to the Acute Fish Toxicity Test: A Focus on Neurotoxicity

**DOI:** 10.1101/2024.09.12.612652

**Authors:** Christoph Schür, Martin Paparella, Christopher Faßbender, Gilly Stoddart, Marco Baity Jesi, Kristin Schirmer

## Abstract

Assessment of potential impacts of chemicals on the environment traditionally involves regulatory standard data requirements for acute aquatic toxicity testing using algae, daphnids and fish (*e.g.*, OECD test guidelines (TG) 201, 202, and 203, respectively), representing different trophic levels. In line with the societal goal to replace or reduce vertebrate animal testing, alternative bioassays were developed to replace testing with fish: the fish cell line RTgill-W1 acute toxicity assay (OECD TG249) and the zebrafish embryo acute toxicity test (zFET, OECD TG236). However, previous studies revealed the lower sensitivity of the RTgill-W1 cell line assay and zFET for some neurotoxic chemicals and allyl alcohol, which is presumably biotransformed in fish to the more toxic acrolein (which is predicted well through the cell line assay). To provide an additional alternative to acute fish toxicity, in this study, we analyzed historic ecotoxicity data for fish and daphnids from the EnviroTox Database. We found a considerable variability in acute fish LC_50_ and acute daphnids EC_50_ values, particularly for neurotoxic chemicals. Comparing sensitivity of these taxonomic groups according to different neurotoxicity classification schemes indicates that fish rarely represent the most sensitive trophic level of the two. Exceptions here most prominently include a few cyclodiene compounds, which are no longer marketed, and a chemical group that could be identified through structural alerts. Moreover, daphnids are more sensitive than fish to acrolein. This analysis highlights the potential of the *Daphnia* acute toxicity test, which is usually a standard regulatory data requirement, in safeguarding the environmental protection level provided by the RTgill-W1 cell line assay and the zFET. This research, rooted in decades of efforts to replace the fish acute toxicity test, shifts the focus from predicting fish toxicity 1-to-1 to emphasizing the protectiveness of alternative methods, paving the way for further eliminating vertebrate tests in environmental toxicology.

## Introduction

Traditional assessment of potential impacts of chemicals on the environment involves standard regulatory data requirements for acute aquatic toxicity testing using algae, daphnids and fish, representing diverse trophic levels. Standardization of experimental setup and conditions in test guidelines (TG), such as those by the Organisation for Economic Co-operation and Development (OECD), does not aim to capture the huge biological variability but to pragmatically enable toxicity comparisons under similar conditions. These include TG 203 for acute fish toxicity and TG 202 on *Daphnia sp.* acute immobilization (as a surrogate for mortality) (OECD, 1992, 2004, 2019).

### Replacing the acute fish toxicity test

Particularly for fish, the motivation for finding adequate replacement methods is strong for ethical, practical, and scientific reasons. Ethically, TG 203 is of considerable animal welfare concern because mortality (or moribundity) is the endpoint. Practically, the relatively high volume of water and chemicals required, the necessary infrastructure, and the 96 hours exposure time are disadvantageous in terms of environmental burden and testing throughput. Scientifically, the basic study design of TG 203 causes uncertainty for experimental variability due to the low number of fish per concentration, the lack of tank replication, absence of an internal positive control and a high degree of flexibility in the test design. It allows for use of eleven different fish species, different endpoints for clinical signs in addition to lethality, and, until its revision in 2019, animal life stages only defined by specimen body length (OECD, 1992, 2019). Moreover, there is considerable uncertainty for the extrapolation of such laboratory data to environmental protection goals (Paparella et al., 2020).

Despite these concerns, the acute fish toxicity test is by far the most frequently conducted aquatic toxicity test using about 50,000 fish per year in Europe, which represents about 50 % of all aquatic vertebrates used for all types of regulatory ecotoxicity testing (Burden et al., 2020; European Commission, 2024). Furthermore, regulatory ecotoxicology faces an ever-growing chemical landscape that was estimated in a 2020 review article to include over 350,000 chemicals and mixtures currently registered on the market worldwide (Wang et al., 2020). For example, the European Union legislation for the Registration, Evaluation, Authorisation and Restriction of Chemicals (REACH) requires the testing of chemicals for acute fish toxicity at a production/import volume above 10 tonnes per year, which currently covers approximately 12,000 chemicals (ECHA, 2024). Therefore, there have been considerable efforts to reduce and replace the use of fish in acute toxicity testing, including the development of a testing strategy (OECD Guidance Document 126 the threshold approach for acute fish toxicity) and the development of alternative methods.

### The threshold approach for acute fish toxicity

The observation that fish are not always the most sensitive test taxon (Weyers et al., 2000) provided the foundation for developing the threshold approach for acute fish toxicity (OECD, 2010). The threshold approach addresses fish toxicity by initially using a single-concentration test (limit test), thus requiring fewer fish compared to the full acute fish toxicity study. The selection of a single concentration is based on the derivation of a threshold concentration (TC) from algae and acute invertebrate (*e.g.*, daphnids) toxicity data. The concept initially described for pharmaceuticals (Hutchinson et al., 2003) was further developed for chemical substances at the European Commission’s Joint Research Centre (Jeram et al., 2005) taking into consideration the requirements of the limit test in OECD TG 203 (European Centre for the Validation of Alternative Methods (ECVAM), 2006). In addition, several publications confirm the potential of the threshold approach to reduce the number of fish used for acute toxicity testing (Hoekzema et al., 2006; Rawlings et al., 2019), also when applied to substances other than industrial chemicals, such as pesticides and pharmaceuticals. Furthermore, retrospective analyses of plant protection products indicate that the overall pain and distress experienced by the fish used in the threshold approach procedure is much less than that of a full acute fish toxicity study, where mortality and adverse symptoms are intended (Creton et al., 2014; OECD, 2002).

### Alternative approaches for acute fish toxicity

Several bioassay-based alternative methods (New Approach Methodologies, NAMs) have been developed or are currently under development to replace the acute fish toxicity test (OECD, 2019) and offer ethical, practical, and economic advantages as well as lower variability, and scientific validation (Paparella et al., 2020). These, most notably, include the fish cytotoxicity test using the cell line RTgill-W1 (OECD TG 249; (OECD, 2021)) and the zebrafish embryo acute toxicity test (zFET; OECD TG 236; (OECD, 2013)). As an additional advantage, data from these alternative methods may support the improvement of computational approaches for predictive ecotoxicology due to lower data variability (Paparella et al., 2020).

However, some limitations pose challenges to the widespread adoption of alternative methods as replacements for the fish acute toxicity test (Westerink, 2013):

1. Studies have highlighted lower sensitivity of the RTgill-W1 cell line assay and zFET to certain modes of neurotoxicity (Glaberman et al., 2017; Klüver et al., 2015; Knöbel et al., 2012; Sobanska et al., 2018; Tanneberger et al., 2013), *i.e.* causing specific adverse functional or structural changes in the nervous system (Legradi et al., 2018). For the RTgill-W1 cell line assay, this is hypothesized to be due to the absence of specific channels or receptors present in nerve but not gill cells (Fischer et al., 2019; Tanneberger et al., 2013). For the zFET, this appears at least to some degree to be related to the oxygen supply of the larvae relying on passive diffusion instead of being dependent on active gill ventilation, as is the case for adults. Active ventilation can be affected through neurotoxicants, rendering older fish life stages more susceptible to respiratory failure syndrome (Kämmer et al., 2024; Klüver et al., 2015).
2. Differences in biotransformation capacity may lead to the underestimation of acute fish toxicity via zFET and the RTgill-W1 cell line assay for compounds more toxic through biotransformation. As prominent example, allyl alcohol is presumably biotransformed in fish to the more toxic chemical acrolein, whereas allyl alcohol is considered to be less toxic in the RTgill-W1 cell line assay and zFET than in the fish acute toxicity test (Glaberman et al., 2017; Klüver et al., 2015; Sobanska et al., 2018). Allyl alcohol is not biotransformed in zebrafish embryos due to the lower expression of the alcohol dehydrogenase isoenzyme adh8a (Braunbeck et al., 2020; Klüver et al., 2014; Knöbel et al., 2012).
3. Another limitation of the zFET is that some molecules may not pass the chorion due to their molecular size, charge and/or structure (> 3000 Da for uncharged molecules; Busquet et al., (2014); Pelka et al., (2017); Braunbeck et al., (2020)).

Currently, an OECD project advancing the threshold approach into a framework of more comprehensive integrated approaches to testing and assessment (IATA) for acute fish toxicity is ongoing (project 2.54 on the <u>OECD Test Guidelines Programme</u>). This IATA will provide adequate information for classification and labelling according to the UN GHS as well as for hazard and risk assessment. It integrates algae and daphnid testing, the zFET, the RTgill-W1 cell line assay, computational approaches, such as quantitative structure activity relationship (QSAR) methods, physico-chemical information, and additional relevant data, *e.g.,* from high-throughput screening and other taxa. However, concerns regarding the ability of the alternative methods to be protective for neurotoxic chemicals have hampered their use for regulatory purposes and have been a motivation for the investigation into whether invertebrate toxicity data is protective for neurotoxicants in fish presented in this study.

### Can invertebrate toxicity data be protective for neurotoxicants in fish?

Recently, it was found that mainly pesticides acting through acetylcholinesterase inhibition and aromatic amines are much more toxic to *Daphnia magna* than to fish (Kienzler et al., 2016). That study analyzed databases available within the OECD QSAR Toolbox 3.2 (Dimitrov et al., 2016; OECD, 2023), exploring extrapolation approaches for avoiding chronic fish testing on the basis of existing data. For example, the neurotoxic insecticide fenitrothion had a medium toxicity to fish but appeared as one of the most chronically toxic chemicals to *D. magna* (Kienzler et al., 2016). In this context, the hypothesis of the data analysis in the present study was to elucidate whether invertebrate toxicity data can be protective for fish for a broad range of neurotoxic compounds. This alternative route to acute fish toxicity prediction could present an important key for the full replacement of acute fish toxicity testing through alternative methods.

In this study, we present a systematic analysis of quality-assured ecotoxicity data from the EnviroTox Database (DB) (Connors et al., 2019) with a focus on chemicals with neurotoxic modes of action. We investigate biological data variability and compare the sensitivity of fish and daphnids for chemicals with a neurotoxic MoA, based on different classification schemes. The resulting insights are contextualized to assess the potential of daphnids to compensate for the described current weaknesses of biological alternative methods for neurotoxicity in the environment.

## Materials & Methods

### Data sources and curation

The data used in this analysis originates from the EnviroTox DB, a curated aquatic toxicity database (Version downloaded January 23^rd^ 2024 from https://EnviroToxdatabase.org). The database was selected because it is compiled based on quality criteria in accordance with the Stepwise Information-Filtering Tool (SIFT; (Beasley et al., 2015)) outlined in Connors et al. (2019) and therefore contains high quality data. A number of filtering criteria were applied to reduce the data to entries that can be reasonably assumed to relate to experiments carried out in accordance with the OECD TG 203 for fish and OECD TG 202 for daphnids.

In short, this includes only data points for acute toxicity testing with the 11 fish species suggested in the TG 203 at 96 h and all daphnid species in the TG 202 at 48 h. For fish, this is limited to outcomes of mortality and moribundity and represented by the LC_50_, Lethal Concentration 50 % or EC_50_, Effective Concentration 50 %. Throughout the paper, LC_50_ is used for fish but encompasses both LC_50_ and EC_50_ data.). For daphnids, the outcome is intoxication/immobilization as a surrogate for mortality (EC_50_). Additionally, if terms within the paper title or effect descriptions were related to egg, embryo or larval data, these data points were excluded to ensure that only tests with juvenile or adult life stages were included. Finally, using the binary filtering option contained in the EnviroTox DB, all entries where the LC_50_ or EC_50_ lay 5-fold above the predicted chemical water solubility were excluded.

To label the tested chemicals as relevant to neurotoxicity, a number of different classification approaches were compared. The EnviroTox DB already includes MoA classifications according to Verhaar, TEST, ASTER, and OASIS (Connors et al., 2019; Kienzler et al., 2017, 2019). TEST and ASTER were deemed to be most informative for the purpose of discerning neurotoxic and non-neurotoxic MoA, due to the inclusion of neurotoxicity-specific classes. These classifications were expanded upon through the inclusion of pesticide-specific modes of action from the Insecticide Resistance Action Committee (IRAC) Mode of Action Database, which is built on experimental data for the purpose of pesticide resistance management (https://irac-online.org/mode-of-action). Additionally, two other sources of MoA categorization were included: I) The candidate compound list by the European Food Safety Authority (EFSA) for the implementation of an *in vitro* test battery focused on developmental neurotoxicity, contributing labeling for 119 chemicals (Masjosthusmann et al., 2020) and II) A list of chemicals that are regularly found in chemical monitoring of European rivers, amounting to over 3,300 compounds with an assigned MoA class (Kramer et al., 2024). The EFSA dataset was augmented by SMILES codes to check overlap with the EnviroTox dataset based on CAS, SMILES, and chemical name. Except for the IRAC categorization, all approaches include both positive and negative compounds to allow for the contextualization of neurotoxically active compounds against chemicals that are not considered neurotoxic or that have not been assigned with a category. Figure 1 gives an overview of the 805 final chemicals included in the analysis (*i.e.*, those for which data for both fish and daphnids in accordance with out filtering criteria are available), their subcategories across classification schemes, and the number of compounds in the different categories. However, the Sankey chart (Figure S1) gives an overview of which subcategories chemicals are assigned to across the different categorization schemes and highlights some discrepancies and limitations of individual approaches, where certain chemicals are labeled neurotoxic in one classification while non-neurotoxic in another. Since the categorization schemes provide a broad range of MoA classifications related to different applications (*e.g.,* pesticides vs. surface water monitoring), categories and chemicals will overlap or differ between these schemes. We see this as a redundancy mechanism strengthening the analysis.

**Figure 1:**
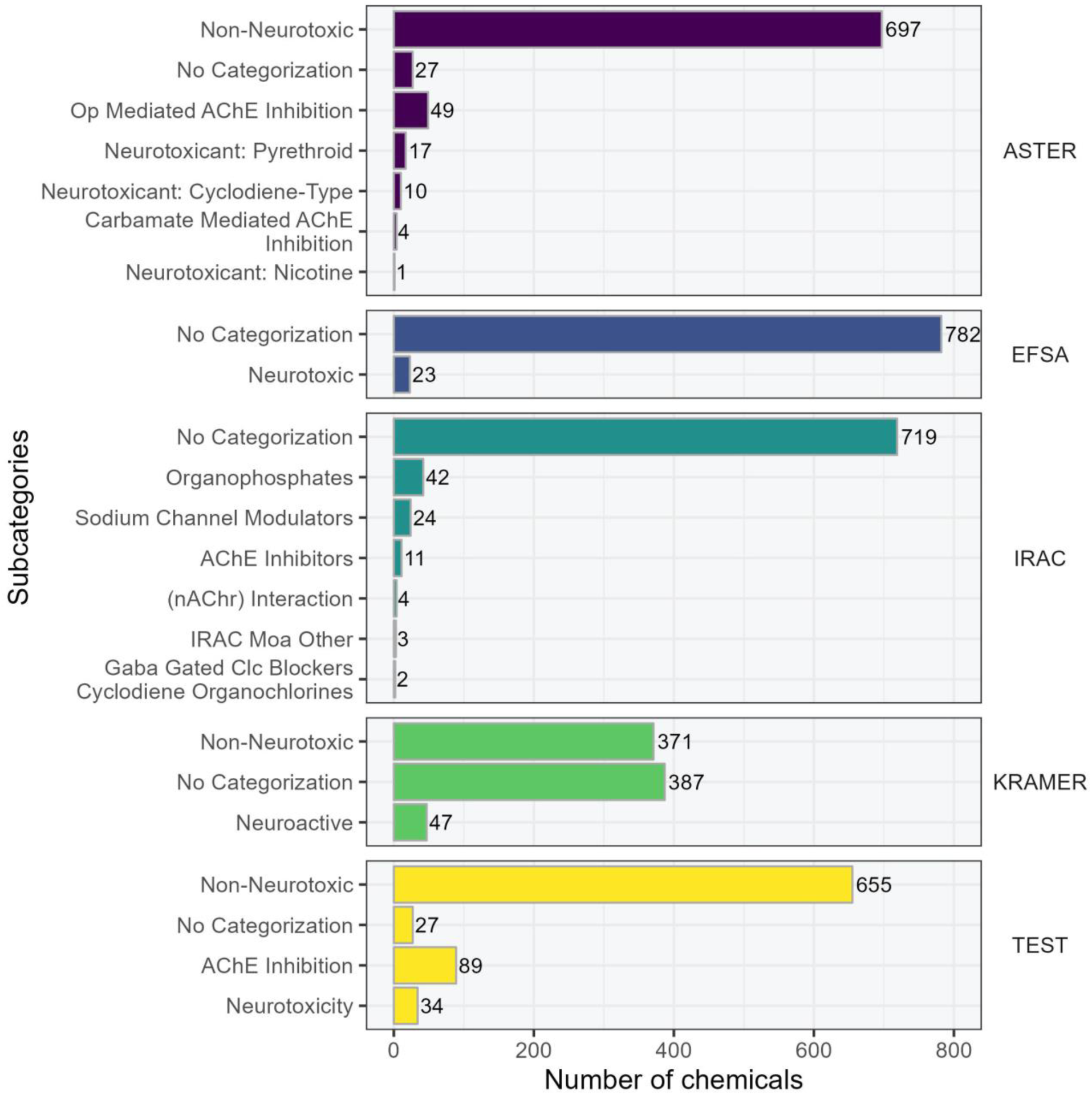
Overview of the 805 chemicals included in the analysis for which both fish and daphnid data in accordance with our filtering criteria was available, their subcategories across classification schemes, and the number of compounds in the different categories of categorization schemes. used to label compounds as potentially neurotoxically active and non-active based on different data sources. Op: Organophosphate; AChE: Acetylcholine Esterase; Clc: Chloride channel; nAChr: Nicotinic acetylcholine receptor.

After merging data from the EnviroTox DB with the supplementary MoA data, the effect values were summarized by calculating median values for each chemical per taxonomic group (fish and daphnids). Afterwards, each of these median values per chemical was matched with each median value for daphnids using the same chemical (based on CAS number). We considered different approaches of summarizing and matching the data and this approach may be justified considering that the current OECD TG allows the use of multiple species and there is no robust knowledge on species-specific sensitivity to the different chemicals on the market. Further reasoning and comparison with alternative approaches for this is described in the supplementary material.

To compare sensitivity between the two taxonomic groups, the ratio R_df_ was calculated according to the following equation (Figure 2):

**Figure 2:**
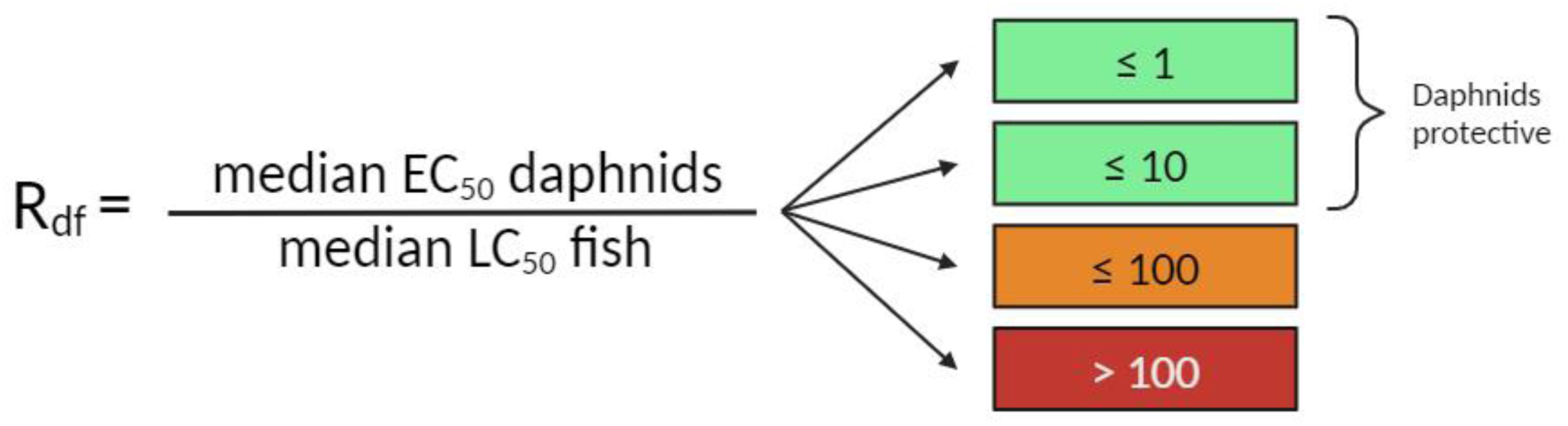
Calculation of R_df_, the ratio between daphnid acute median EC_50_ and fish acute median LC_50_. The colored bars to the right indicate different levels of concern: R_df_ ≤ 1 and ≤ 10 are considered unproblematic, since they indicate that daphnids are either more sensitive or up to a factor of 10 less sensitive than fish to the respective compound and can, thus, be considered protective for fish. R_df_ values above 10 are critical, since they indicate that fish are more sensitive than daphnids by at least a factor of 10. The figure was created with BioRender.com.

This ratio gives an intuitive understanding of relative sensitivity between the two taxonomic groups. A ratio of 1 indicates that the two groups are equally sensitive to the respective chemical, while values below 1 relate to a higher sensitivity in daphnids. For the analysis, ratios above 10 were considered to be potentially problematic, because they indicate that, for the respective chemical, fish are at least 10 times more sensitive than daphnids in the standardized test guideline tests.

### Data analysis

Data transformation, analysis, and visualization was carried out using R (R Core Team, 2018) with RStudio (V2023.12.1+402) (RStudio Team, 2024) and the *tidyverse* package (Wickham et al., 2019). Ridgeline plots were created using the *ggridges* package (Wilke, 2024). Figure 2 was created with BioRender.com. The Sankey chart (Figure S1) was created with *ggsankey* (Sjoberg, 2024). Random subsampling of data points for Figure 3 was done using the slice_sample function from the *dplyr* package. Kruskal-Wallis rank sum test was performed using the *kruskal.test* function of the *stats* package. SMILES codes for the supplementary data were retrieved using the python package *cirpy* (https://cirpy.readthedocs.io). The layout of Figure 5 was adjusted using Inkscape 1.2.2 (732a01da63, 2022-12-09) for Windows. The R code used to produce the analyses and figures in this publication is available on GitHub (### link currently removed to anonymize the manuscript ###).

**Figure 3:**
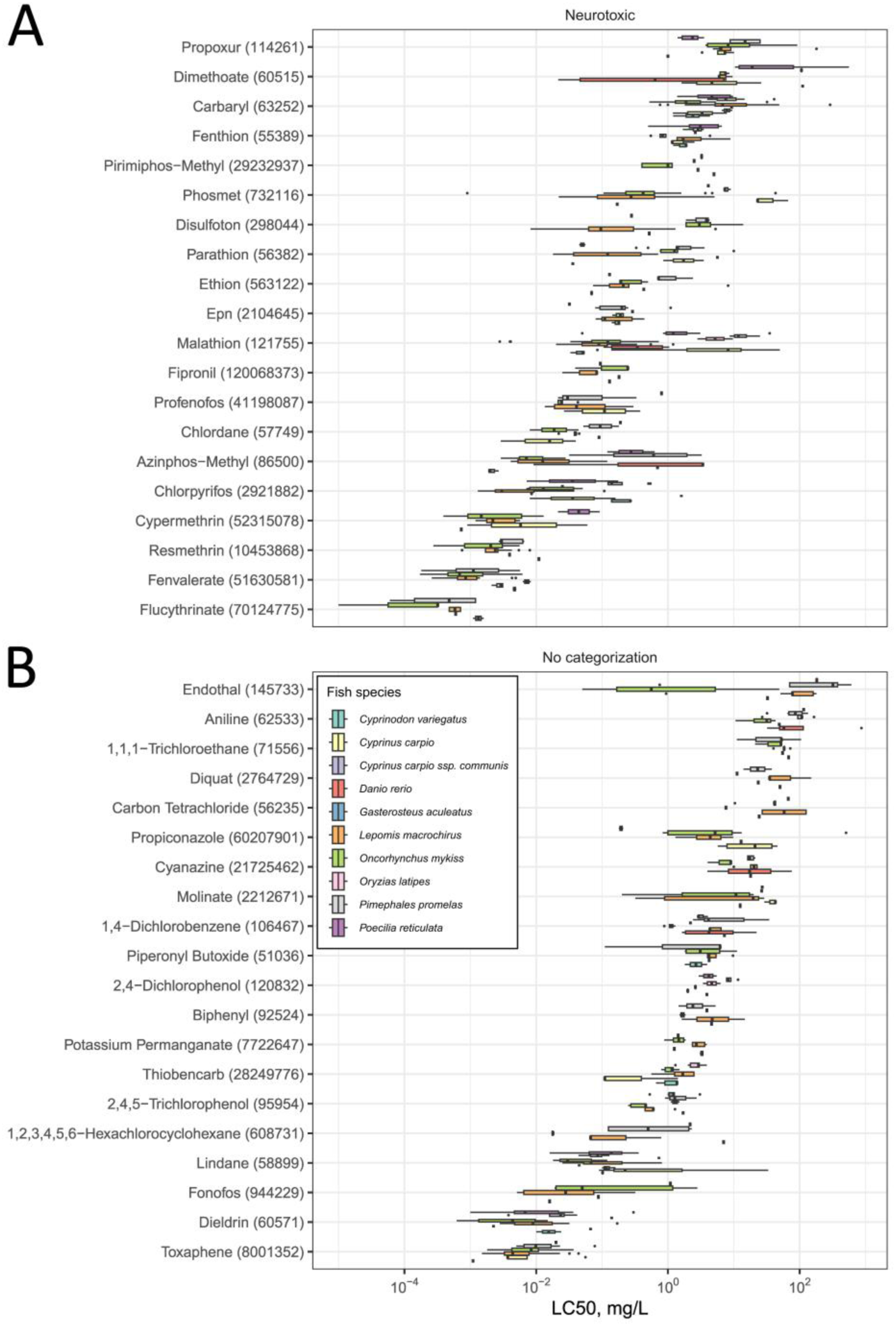
Variability of LC_50_ values across more than four species in fish for 20 randomly selected chemicals from the chemicals assigned (A) a neurotoxic MoA (32 chemicals total) and (B) no categorization (90 chemicals total) through IRAC. CAS numbers of test chemicals are included in brackets. Figure S5 displays significance levels of species variability across the whole chemical set.

## Results & Discussion

### Overview of the data and variability across taxonomic groups

The analysis was limited to 805 chemicals for which acute toxicity data on both fish and daphnids (related to OECD TG 203 and 202, respectively) were contained in the EnviroTox DB, and for which MoA classifications were available. Tables S1 and S2 give an overview of effect types, test statistics, and different species included in the dataset. As an initial step, we characterized this dataset by visualizing the distribution of median EC_50_/LC_50_ values across the two taxonomic groups fish and daphnids (Figure S2, A) as well as the inter-species variability of the data for the two taxonomic groups (Figure S2, B+C; Figure S2, D contains a similar characterization of amphibian toxicity data). The range of median values is similar across fish and daphnids with slightly lower values for the latter, indicating a generally higher sensitivity. This is also indicated by the quartiles (vertical lines in the plots A-C in Figure S2) being lower as well (*i.e.*, towards lower effective concentrations and, hence, higher sensitivity) compared to the quartiles in the fish distribution.

Inter-species variability for the selected compounds seems to be influenced by data availability as can be seen for non-summarized data points (*i.e.,* raw data points, not median values) in Figure S2 B-D. The fish species with the most data available after pre-filtering (indicated by the numbers given in the y-axis labels in Figure S2 B-D) appear to be approximately normally distributed and largely overlap in their range, while species with fewer tested chemicals seem to have more unevenly distributed LC_50_ values (Figure S2, B). This also holds true for the daphnid data (Figure S2, C), albeit this subset is, for the most part, dominated by a single species (*Daphnia magna*). *Daphnia carinata* and *Daphnia laevis* generally appear to be more sensitive or have been tested on more-toxic chemicals with their distributions centered on the left of the scale (lower EC_50_ values, higher toxicity). Contrary to that, data for *Daphnia longispina* and *Daphnia obtusa* are less evenly distributed, which likely is related to the low number of data points (5 and 6, respectively). Accordingly, observations based on such low numbers of data points should be taken *cum grano salis*. There is a poor correlation between the number of data points per chemical and the coefficient of variation (CV; R^2^ of 0.11 and 0.29 for daphnids and fish, respectively; Figure S3). The CV is the relative standard deviation, *i.e.*, the standard deviation divided by the mean, expressed as percentage. Here, it supports the notion that the CV does not necessarily increase with more data points, but it becomes more robust (Figure S3).

From the overlap of the two datasets, the ratios of median daphnid effect values divided by the median fish effect values were calculated as a marker of sensitivity differences between the two taxonomic groups (Figure 2, Figure S2, E). At this point, the analysis is agnostic about the associated MoA. The distribution is skewed towards the left of a ratio of one, *i.e.*, higher sensitivity for daphnids with only a small fraction of ratios indicating a higher sensitivity of fish by a factor of 10 or 100 compared to daphnids (orange and red area, respectively). A similar pattern emerges when corresponding median fish LC_50_ and daphnids EC_50_ values are plotted against each other to visualize the correlation differently (Figure S4). Here, it can be observed that over all 805 chemicals, daphnids are the more sensitive taxonomic group and only for about 4 % of the chemicals fish are more than 10-fold more sensitive than daphnids.

### Biological data variability

Since the analysis is based on ecotoxicological *in vivo* data, it is important to acknowledge the high biological variability associated with such information. Here, we restrict to chemicals that are covered by the IRAC categorization scheme. 32 and 90 chemicals had been tested on daphnids and at least on four fish species and had a neurotoxicity MoA or no categorization in the IRAC categorization scheme, respectively. We use IRAC as the reference categorization here because we expect that knowing the MoA of a pesticide constituent gives the highest reliability of a correct categorization as opposed to, *e.g.*, a purely predicted MoA. This assumption comes with the caveat that IRAC only contains 86 active compounds and is biased towards insecticides. To keep the two groups comparable, we randomly subsampled 20 chemicals that have been tested on at least four different fish species from each of those two groups. We visually oppose them in Figure 3, displaying the range of LC_50_. Even though data were selected for only one experimental duration and limited as closely as possible to OECD TG 203 conformity and excluded LC_50_ values which were more than 5-fold above the chemicals water solubility limits, we still observed high variability of the experimental outcomes. In some cases, the LC_50_ values for a single species and chemical span several orders of magnitude. Visually, this is more distinct for the neurotoxic compounds in Figure 3, A. To confirm this statistically we performed a Kruskal-Wallis rank sum test between the species for each of the chemicals from the original pool of 32 and 90 neurotoxic and uncategorized chemicals (Figure S5). Here, the number of significantly different (p < 0.05) distributions of LC_50_ values is far higher for the chemicals with an assigned neurotoxic MoA according to IRAC than for chemicals without an IRAC MoA (84.4 % vs. 44.4 % across 32 and 90 chemicals, respectively; Figure S5).

A figure similar to the boxplot in Figure 3 for daphnids (at least two tested species) is included in the supplementary material as Figure S6. There, the data mostly comes from studies using *D. magna* as the primary test species (see Table S2), which hinders the comparison of inter- and intra-species variability. Additionally, we visualize the CV, *i.e.,* the standard deviation (SD) normalized against the mean, across the broad categories of non-neurotoxic, no categorization, and neurotoxic across all chemicals and categorization schemes (Figure 4, A). This is done for fish data, but limited to the chemicals that have been tested on both fish and daphnids. Notably, since this is based on all categorization schemes, chemicals can be assigned to several categories at once (see Figure S1), leading to duplicate data points. The median of the CVs for neurotoxic chemicals is about 100 %, which is about twice the median of the CVs of other chemicals. However, the lower quartile of the neurotoxic CVs is overlapping with the upper quartile of the other chemicals CVs. As a refinement, Figure 4B focuses only on IRAC-labeled chemicals. Here, the median of the CVs is also higher in the group of chemicals that have been assigned a neurotoxic MoA compared to those without an IRAC classification. In conclusion, toxicity outcomes in fish appear more variable for chemicals with an assigned neurotoxic MoA. Consequently, when only a few (pseudo-)replicates are available, this higher uncertainty about the “true” median LC_50_ could influence the R_df_ values derived from such data, potentially leading to very high or very low R_df_ values.

**Figure 4:**
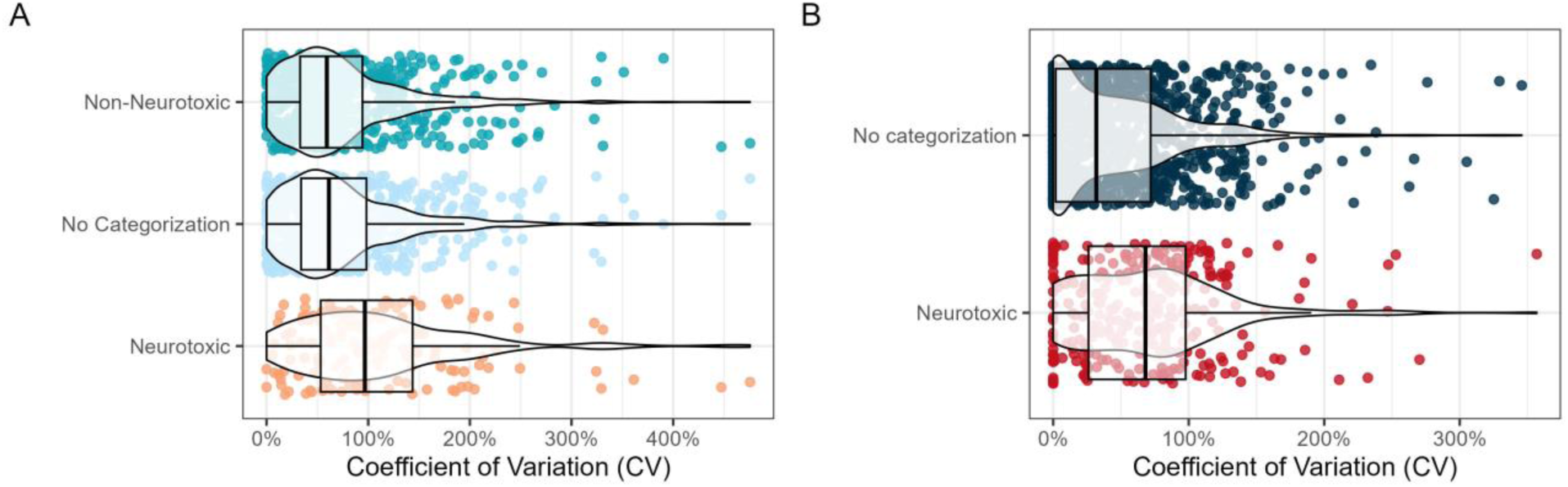
Coefficient of variation across fish for all the chemicals in the dataset that also have been tested on daphnids. (A) assigned to the broad classes of non-neurotoxic, no categorization, and neurotoxic across all five categorization schemes and (B) only the chemicals with an assigned MoA in the IRAC categorization scheme.

### Neurotoxicity classification schemes

Since the main goal of this study was to analyze whether daphnid acute data could be protective of fish acute mortality for neurotoxic compounds, it was important to establish a ground truth for the MoA of the chemicals in the dataset. In other terms, it was necessary to introduce knowledge on which chemicals are considered to cause neurotoxicity. For this, six classification schemes were selected that, for the most part, contained positive (*i.e.*, potentially neurotoxic), as well as non-neurotoxic compounds (Figure 1). For the sake of consistency, subcategories for neurotoxicity were retained where available (some category names were adapted for clarity), but data for chemicals that were either labeled as something other than neurotoxic (“Non-Neurotoxic”) or that were not labeled in the specific scheme at all (“No Categorization”) were summarized separately.

Within this analysis we considered that chemical median ratios between daphnids EC_50_ and fish LC_50_ values of less than 10 (*i.e.* fish are not more than 10 times more sensitive than daphnids) are in agreement with the hypothesis that daphnid EC_50_ values are sufficiently protective in the absence of fish LC_50_ values. This consideration is based on several observations:

1. Chemical-specific LC_50_ and EC_50_ values for fish and daphnids generated according to OECD TG may range several orders of magnitude with a typical CV of 100 % for chemicals with neurotoxic MoA, which is about twice as high as for non-neurotoxic chemicals (see section above). Irrespective of MoA, this observation is in line with the findings of Hrovat et al. (2009), who found variability of up to six orders of magnitude for fish LC_50_ derived from the ECOTOX database, albeit without selecting for specific experimental setups. Another analysis by Braunbeck et al. (2020) applied more stringent filtering to manually curated fish acute toxicity test data, which resulted in a smaller dataset of 58 chemicals, with only two chemicals with neurotoxic MoAs in accordance with our classifications. They kindly made the raw data available to us, enabling a comparative analysis with consistent data handling and metric calculation. We found that only 18 of the chemicals in their dataset are also part of the current analysis, albeit none of them are classified as neurotoxic. Furthermore, in that analysis, essentially for non-neurotoxic chemicals, CVs ranged up to 150 % with a median of about 20 % (supplementary Figure S6). Fischer et al. (2019) (see Figure 5 therein) supports the particularly high fish inter-species variability for the neurotoxic pesticide malathion (> 4 orders of magnitude). Scholz et al. (2016) reported variability of fish LC_50_ values of more than 50-fold for chemicals that are toxic mainly after bioactivation. Paparella *et al*. (2020) further summarized the current knowledge on variability of data related to OECD TG 203.
2. Within current regulatory practice, safety factors of 1000 are applied to acute aquatic toxicity data for the derivation of Predicted No Effect Concentrations (PNEC). This approach aims to arrive at protective values for the environment based on acute laboratory experiments. In light of the high biological variability we (and others) observe within highly standardized experimental designs, we argue that this may not adequately translate to protective PNECs.
3. UN GHS classification for aquatic toxicity stratifies acute toxicity into 10-fold ranges, which are then used to derive M-factors for the purpose of toxicity-based weighting of components for mixture assessment. Similarly to the argument above, these ranges may be too granular to adequately account for the observed biological variability.

### Median ratios across MoA categories

Generally, the emerging patterns indicate that for most neurotoxic compounds, daphnids are more sensitive than fish compared to non-neurotoxic compounds (Figure 5). Exception are the “*Cyclodiene-type*” neurotoxins in ASTER and the analogous “*Gaba-gated Clc Blockers Cyclodiene Organochlorines*” in IRAC. The cyclodiene-type category in ASTER contains four chemicals with an R_df_ above 10, namely, dieldrin, endosulfan, endosulfan sulfate, and endrin. These chemicals also appear within the EFSA, Kramer and TEST classifications, which do not specify cyclodiene-type neurotoxicity as such (blue triangles visualize these chemicals across the categories in Figure 5B). In addition, two pyrethroids, *i.e.,* flucythrinate and resmethrin have a ratio slightly above 10 (13.8 and 19.7, respectively) within the ASTER pyrethroid category as well as within the other classification systems.

**Figure 5:**
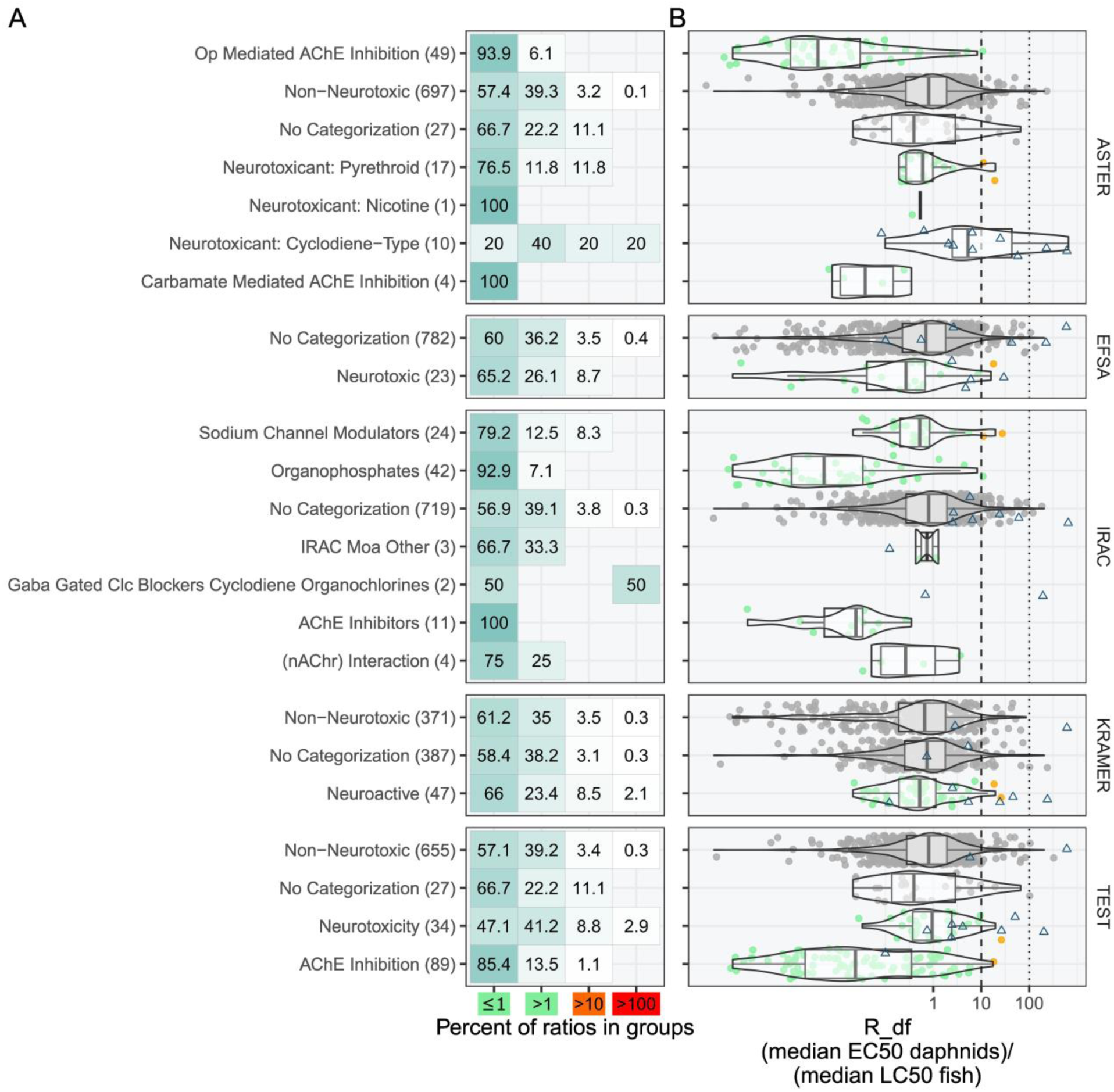
A: Heatmap/table of median ratios across the classification schemes and subclasses. Median ratios are divided into the groups ≤1, between >1 and ≤ 10, between >10 and ≤100, and >100. Darker shading of cells correlates to higher percentages. B: Violin plot of the ratio distributions across the different categorization schemes and subcategories. Grey dots are not relevant to neurotoxicity. Colorful dots indicate the range of R_df_ ratios for neurotoxicity-related endpoints (green ≤ 10; orange 10-100). Blue triangles indicate chemicals that are part of the ASTER subcategory “Neurotoxicant: Cyclodiene-Type”). Op: Organophosphate; AChE: Acetylcholine Esterase; Clc: Chloride channel; nAChr: Nicotinic acetylcholine receptor.

Out of the 32 compounds with ratios ≥10, seven were categorized as neurotoxic in at least one of the classification schemes (five out of these seven chemicals were labeled as neurotoxic by at least three out of five classification schemes). The observation of most compounds being labeled as neurotoxic in several classification schemes strengthens confidence that the approach applied in this analysis is highly reliable through redundancy. All the five cyclodienes ratios with > 10 (dieldrin, endosulfan, endosulfan sulfate, endrin, 1,2,3,4,5,6-hexachlorocyclohexane (lindane)) are covered in Annex A of the Stockholm convention. The two pyrethroids (flucythrinate and resmethrin) are excluded from being used as ingredients in plant protection products through the European Commission Regulation (EC) No 2076/2002 (Commission Regulation (EC) No 2076/2002 of 20 November 2002 Extending the Time Period Referred to in Article 8(2) of Council Directive 91/414/EEC and Concerning the Non-Inclusion of Certain Active Substances in Annex I to That Directive and the Withdrawal of Authorisations for Plant Protection Products Containing These Substances (Text with EEA Relevance), 2002). Hence, all compounds labeled as neurotoxic that have a median ratio higher than 10 are no longer relevant to the market. This is true for the European market for the REACH-regulated chemicals and globally for those covered by the Stockholm convention ratified by 186 countries (https://treaties.un.org/Pages/ViewDetails.aspx?src=IND&mtdsg_no=XXVII-15&chapter=27&clang=_en; Table 1). Resmethrin products are no longer sold or distributed across the United States since 2015 and the use of remaining product is highly limited (https://www.epa.gov/mosquitocontrol/permethrin-resmethrin-d-phenothrin-sumithrinr-synthetic-pyrethroids-mosquito). It needs to be stated, however, that the CVs for fish LC_50_ and daphnids EC_50_ values are particularly high for these chemicals (Table 1), such that the daphnids/fish median ratio contains considerable uncertainty.

**Table 1:**
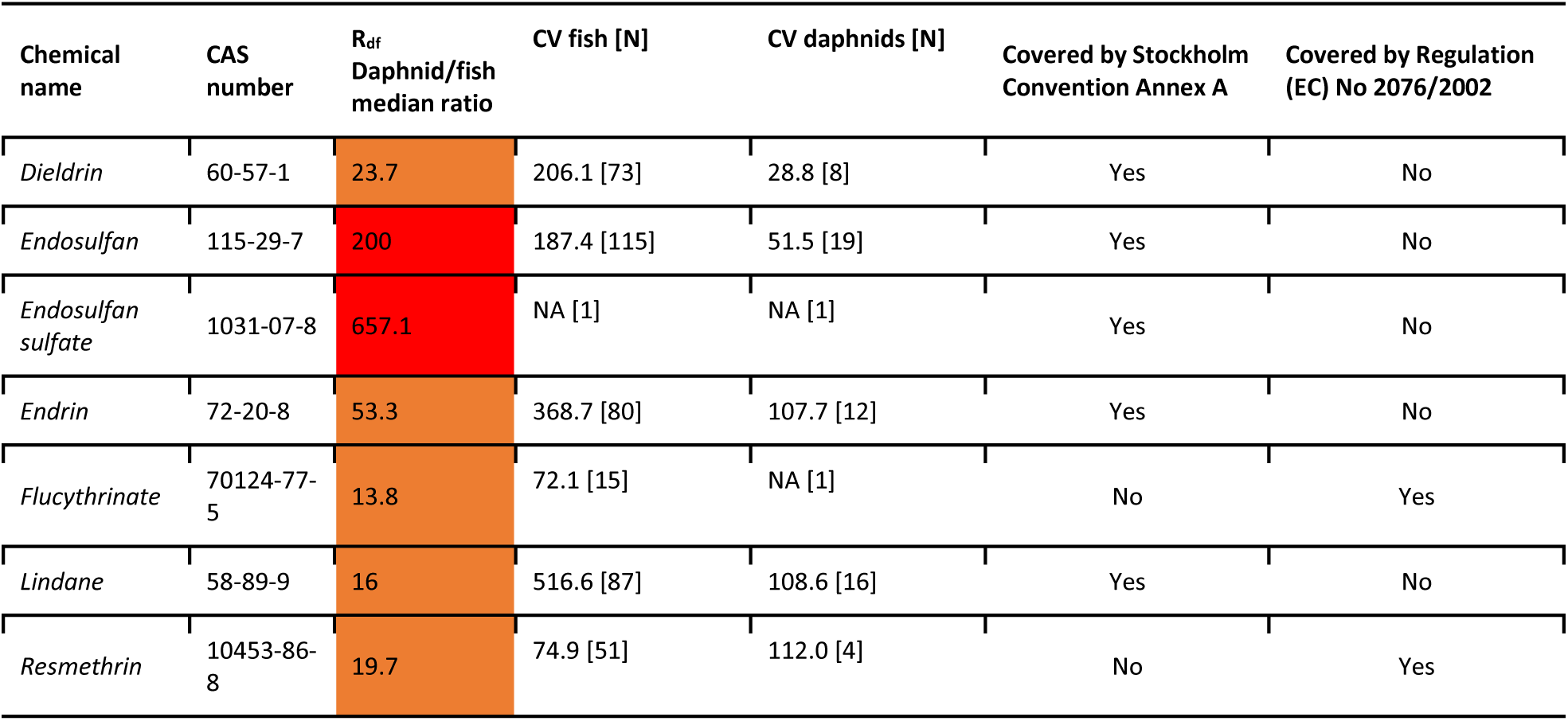
Overview of the seven chemicals labeled with a neurotoxic MoA in at least one of the classification schemes and with a median ratio above 10. Five out of seven compounds are covered in the “Elimination” Annex of the Stockholm convention, while the other two are prohibited from use in plant protection products through EC regulation No 2076/2002. CV = Coefficient of variation.

Moreover, beyond neurotoxicity, just 3.9 % of the 805 chemicals for which both fish and daphnid data were available showed a median R_df_ above 10 (Table S3 in the supplementary material), which supports that within the current overall chemical universe, fish LC_50_ values rarely drive PNEC derivation and classification. Importantly, the 3-4% of non-neurotoxic chemicals with ratios higher than 10 are of low concern, if alternative methods to the fish test are used (TG 236 and TG 249) because, based on current knowledge, they are protective for chemicals with non-neurotoxic mechanisms.

In our presentation, the comparison of daphnids and fish toxicity data is based on the calculation of the chemicals’ median LC_50_ or EC_50_ values averaging all LC_50_ or EC_50_ values within the trophic level. It is acknowledged that also different approaches may be used, *i.e.*, grouping by chemical and species or no grouping and matching all fish LC_50_ values with all daphnids EC_50_ values. The arguments and results for these alternative approaches are provided in the supplemental information under “Comparison of different data matching approaches”. Importantly, the overall pattern emerging from these other approaches is similar (Figures S9 and S10, Table S4).

### Chemicals underestimated in the fish cell acute toxicity assay and zFET

In terms of NAMs, assays completely rendering the use of animals (vertebrates and invertebrates alike) obsolete are preferable. Accordingly, this section focuses on the RTgill-W1 cell line assay and chemicals that were reported as underpredicted. Tanneberger et al. (2013) analyzed the correlation between EC_50_ from the fish cell acute toxicity assay and fish acute LC_50_ values. Out of the 35 compounds investigated in that study, five compounds had fish LC_50_ values at least 10 times lower than the EC_50_ derived from the *in vitro* experiments: permethrin (190-fold), caffeine (18-fold), lindane (63-fold) (all three neurotoxicants), allyl alcohol (2700-fold), and 4-fluoroaniline (12-fold). Accordingly, the authors of that study, based on which the OECD TG 249 (OECD, 2021) was later developed, found these compounds to be underpredicted by the RTgill-W1 cell line assay. Those compounds were specifically sought out in our dataset (where available) and are summarized in Table 2. Since data was not always available at the desired time point of 48 h for daphnids, the filter criteria had to be adapted to be less stringent. Expanding towards daphnid experiments performed at 24 h (in addition to 48 h) and LC_50_/mortality data in addition to EC_50_ and immobilization, allowed four of the chemicals of interest (permethrin, lindane, caffeine and allyl alcohol) to be included. No daphnid data was available for 4-Fluoroaniline.

**Table 2:** Outlier chemicals that Tanneberger *et al*. (2013) found to be underpredicted by fish cell acute toxicity assays compared to fish and their median LC_50_/EC50 in fish and daphnids in our analysis as well as median ratios, coefficient of variation (CV) and number of data points [N]. NA = not applicable.

| Chemical name | CAS number | R<sub>df</sub> Daphnid/fish median ratio | Median fish | CV fish [N] | Median daphnids | CV daphnids [N] |
| --- | --- | --- | --- | --- | --- | --- |
| *Allyl alcohol* | 107-18-6 | 0.4 | 0.59 | 57.3 [9] | 0.25 | NA [1] |
| *Permethrin* | 52645-53-1 | 0.2 | 0.01 | 87.7 [109] | 0.001 | 197.8 [42] |
| *Caffeine* | 58-08-2 | 2.8 | 151 | NA [1] | 422 | 87.8 [2] |
| *Lindane* | 58-89-9 | 20.8 | 0.09 | 516.6 [87] | 1.79 | 116.6 [40] |

Within this analysis, the R_df_ are in an unproblematic range (< 3) for allyl-alcohol, permethrin and caffeine, while for lindane the ratio of about 21 is only slightly above the ratio of 10 and needs to be contextualized with its fish CV of 517 %, rendering this ratio uncertain.

Importantly, this analysis puts allyl-alcohol in a different light. Rather than being an example for a compound considered “difficult” for alternative testing schemes due to its presumed toxification via biotransformation in fish, it is here considered an example of outliers in the performance of alternative methods that may be well covered with the obligatory daphnid test.

In summary, the toxicity of the outliers not sufficiently covered by the RTgill-W1 cell line assay is covered by daphnid data, thus reducing an uncertainty in the toolbox of bioassay-based NAMs for fish acute toxicity.

In a similar analysis, comparing zFET data with fathead minnow (*Pimephales promelas*) acute toxicity data, Knöbel et al. (2012) identified four chemicals for which zebrafish embryos were more than ten times more sensitive: 2,3-dimethyl-1,3-butadiene (19-fold less toxic), rotenone (15-fold), allyl alcohol (1059-fold), and acrolein (22-fold). With the exception of allyl alcohol (which is covered by daphnids with an R_df_ of 0.4, see Table 2), these are predicted well through the RTgill-W1 cell line assay (Tanneberger et al., 2013). Rotenone likewise is covered well by the daphnids (R_df_ of 0.23), while not data is available for 2,3-dimethyl-1,3-butadiene.

### Complementing fish acute toxicity alternative methods

Given the currently available data, TG 236 and 249 are considered to be of limited use for chemicals with a neurotoxic MoA. In the still only recently (in 2021) instated TG 249, this is acknowledged to concern chemicals specifically acting on ion channels or receptors typical for brain tissues (OECD, 2021). Given the other underpredicted chemical, allyl alcohol, the authors argue that the assay needs to be conducted on both the original chemical as well as relevant transformation products, should indications arise that those transformation products contribute to or drive toxicity. This is based on the observation that the more toxic transformation product acrolein was predicted well through the RTgill-W1 assay.

Complementarily to that, our analysis demonstrates that daphnid acute toxicity data from studies conducted according to OECD TG 202 can be considered protective for fish for such compounds due to the higher sensitivity of daphnids. Exceptions, as we have shown, are seven out of 141 compounds with neurotoxic mechanisms, mainly belonging to a distinct group of chemicals. They were identified and predicted to be neurotoxic via the ASTER tool and the IRAC database, showcasing how computational tools and biological assays can further complement each other. The chemicals of concern are included either in the Annex A (“Elimination”) of the Stockholm convention or the EC regulation No 2076/2002 and, accordingly, not relevant for markets in the European Union and in the 186 countries that ratified the Stockholm convention.

### Limitations and outlook

The determination of the ground truth for neurotoxic MoAs, *i.e*., the way in which chemicals considered to cause neurotoxicity were identified, presents an inherent bias in this analysis (see the section on neurotoxicity classification schemes). Therefore, a broad range of classification schemes was selected, spanning both very specific pesticide MoAs (*e.g.*, IRAC) as well as a broad selection of chemicals relevant for freshwater monitoring (*e.g.*, the data from Kramer *et al*. (2024)). The fact that the neurotoxic chemicals with a median ratio above 10 were for the most part flagged by multiple classification schemes shows that this redundancy mechanism added robustness to the analysis and demonstrates that its findings are indeed not biased. However, it is possible and likely that new chemicals that are more toxic to fish than to daphnids may emerge in the future, probably belonging to completely new chemical classes. Therefore, this analysis serves as an example that the current regulatory framework can benefit from the modular adoption of alternative methods. For this, a concerted effort is required towards full integration of such methods into a new framework that accounts for the strengths and weaknesses of each alternative method as opposed to expecting a single method to serve as a full one-to-one replacement of a traditional animal test. Even though our analysis focuses on data derived from biological toxicity assays, we acknowledge the high potential of *in silico* methods, such as QSARs with and without the use of machine learning (Gasser et al., 2024; Kleinstreuer & Hartung, 2024; Muratov et al., 2020). They may serve as the uniting puzzle piece that enables the integration of different kinds of data (physicochemical, *in vivo*, *in vitro*, *etc.)* into models that ultimately can allow for predictions across a broad space of endpoints and biology, given the availability of sufficiently high-quality data.

However, this work already supports a fully NAM-based IATA for fish acute toxicity including the zFET, the RTgill-W1 cell line assay and computational approaches, in addition to daphnids and algae testing, supporting a full replacement of the acute fish toxicity test.

Finally, considering the observed variability of acute fish and daphnids data generated according to OECD TG, it is scientifically warranted to discuss how the UN GHS approach could be improved. The current approach foresees a deterministic classification of chemicals into the acute aquatic 1 category or no classification, depending on the lowest LC/EC_50_ value from fish, daphnids, and algae being below or above 1 mg/L. However, considering the confirmation within the present analysis that LC/EC_50_ values from fish and daphnids may span several orders of magnitude, an approach providing concentration ranges rather than point estimates may be more adequate. Similarly, reflection is needed on the current UN GHS stratification of aquatic LC/EC_50_ values into 10-fold ranges, which are then used to derive M-factors for the purpose of mixture assessment.

### Towards Next Generation Hazard and Risk Assessment

In the context of a chemical hazard assessment that is based on the 3Rs principle (replacement, reduction, and refinement of animal tests), this analysis showcases how the integration of alternative systems can benefit the replacement of vertebrate species. Since daphnid toxicity data is currently a standard regulatory data requirement, testing on daphnids is conducted irrespective of whether fish are tested as well. Therefore, focusing on a combination of alternative methods in conjunction with the anyway created daphnid data leads to a net reduction in testing on animals, since fish tests can be omitted. Interestingly, acute daphnid EC_50_ values appear also protective for acute amphibian EC_50_ values, irrespective of the underlying MoA and as far as available within the EnviroTox DB (see Figure S8). A next generation risk assessment (NGRA) framework that is fit for the challenge of an ever-growing chemical landscape and concerned with environmental health foremost needs to be considerate of the protection goals. Achieving high capacities of chemical testing while ensuring environmental protection will likely be feasible sooner through the combination of alternative methods with traditional daphnid testing. This way of considering strengths and shortcomings of individual methods creates a framework that covers a range of outcomes rather than aiming at replacing highly variable acute fish toxicity testing 1-to-1.

Nonetheless, we consider the use of daphnids as safeguards an intermediate solution that does not negate the need for the further development and adoption of approaches that are completely free of animal use. Here, the already more advanced fully non-animal based NGRA concepts for human health may stimulate the evolution of environmental NGRA concepts (Langan et al., 2023). Referring to the approach taken here for analyzing mechanism of action, this work presents also an important stepping-stone towards such a broader, fully animal-free, IATA for environmental safety.

## Conflicts of interest

The authors declare no conflicts of interest.

## Acknowledgements

This work was made possible through the Swiss data science center (SDSC) grant “Enhancing Toxicological Testing through Machine Learning” (project No C20-04) and partly carried out in the framework of the European Partnership for the Assessment of Risks from Chemicals (PARC) and has received funding from the European Union’s Horizon Europe research and innovation program under Grant Agreement No 101057014. The work of MP at the Medical University is co-financed *via* PARC and the Austrian Federal Ministry for Climate Action, Environment, Energy, Mobility, Innovation and Technology, Department V/5— Chemicals Policy and Biocides. We thank Thomas Braunbeck (University of Heidelberg, Germany) for providing us with the raw data from his analysis of fish LC_50_ variability (Braunbeck et al. 2020), which allowed us to compare our analysis.

## Author contribution

CS: Conceptualization, Data curation, Formal Analysis, Investigation, Methodology, Software, Validation, Visualization, Writing – original draft

MP: Conceptualization, Data curation, Formal Analysis, Funding acquisition, Investigation, Methodology, Software, Validation, Visualization, Writing – original draft

CF: Conceptualization, Writing – original draft, Writing – review & editing

GS: Conceptualization, Writing – review & editing

MBJ: Conceptualization, Project administration, Supervision, Funding acquisition, Writing – review & editing

KS: Conceptualization, Project administration, Supervision, Funding acquisition, Writing – review & editing

## Notes

### Competing Interest Statement

The authors have declared no competing interest.

