## Supplementary Material for "Daphnids Can Safeguard the Use of Alternative Bioassays to the Acute Fish Toxicity Test: A Focus on Neurotoxicity"

**Table S1: Overview of the number of data points (and percentages) across trophic levels, effect types, and test statistics. These are not summarized, hence can include multiple values per experiment, that later are summarized by CAS number.**

| Trophic level | Effect type | Experimental duration (h) | Test statistic | N | Total N per trophic level | Percentage |
|---|---|---|---|---|---|---|
| Fish | Mortality | 96 | $LC_{50}$ | 11379 | 13552 | 84 % |
| | Immobilization | 96 | $LC_{50}$ | 2 | 13552 | < 1 % |
| | Mortality | 96 | $EC_{50}$ | 2131 | 13552 | 15.7 % |
| | Immobilization | 96 | $EC_{50}$ | 40 | 13552 | < 1 % |
| Invertebrate | Immobilization | 48 | $EC_{50}$ | 2885 | 2885 | 100 % |

**Table S2: Number of data points per species relevant to the OECD TG for fish acute toxicity and daphnia acute mortality (TG 203 and TG 202).**

| Trophic level | Species Latin name | N | Percentage |
|---|---|---|---|
| Fish | *Oncorhynchus mykiss* | 4325 | 31.9 |
| | *Lepomis macrochirus* | 3740 | 27.6 |
| | *Pimephales promelas* | 3344 | 24.7 |
| | *Poecilia reticulata* | 572 | 4.2 |
| | *Oryzias latipes* | 429 | 3.2 |
| | *Cyprinus carpio* | 417 | 3.1 |
| | *Cyprinodon variegatus* | 365 | 2.7 |
| | *Danio rerio* | 306 | 2.3 |
| | *Gasterosteus aculeatus* | 35 | 0.3 |

| Trophic level | Species Latin name | N | Percentage |
|---|---|---|---|
| | *Cyprinus carpio ssp. Communis* | 14 | 0.1 |
| | *Dicentrarchus labrax* | 1 | 0 |
| | *Pagrus major* | 4 | 0 |
| Invertebrate | *Daphnia magna* | 2367 | 82 |
| | *Daphnia pulex* | 324 | 11.2 |
| | *Ceriodaphnia dubia* | 123 | 4.3 |
| | *Daphnia carinata* | 25 | 0.9 |
| | *Daphnia sp.* | 13 | 0.5 |
| | *Daphnia laevis* | 9 | 0.3 |
| | *Daphnia longispina* | 7 | 0.2 |
| | *Daphnia obtusa* | 6 | 0.2 |
| | *Daphnia spinulata* | 6 | 0.2 |
| | *Ceriodaphnia lacustris* | 2 | 0.1 |
| | *Moinodaphnia macleayi* | 2 | 0.1 |
| | *Daphnia galeata* | 1 | 0 |

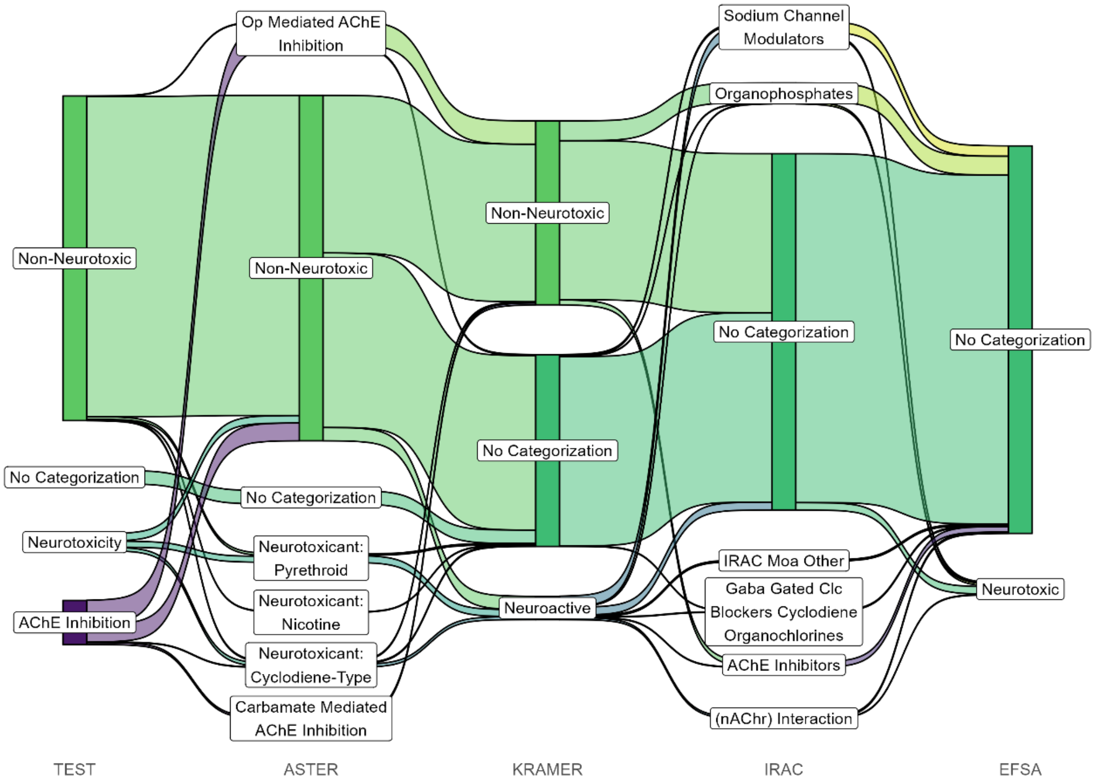


**Figure S1: Sankey diagram of the distribution of chemicals across subcategories in different categorization schemes. (**Op: Organophosphate; AChE: Acetylcholine Esterase; Clc: Chloride channel; nAChr: Nicotinic acetylcholine receptor)

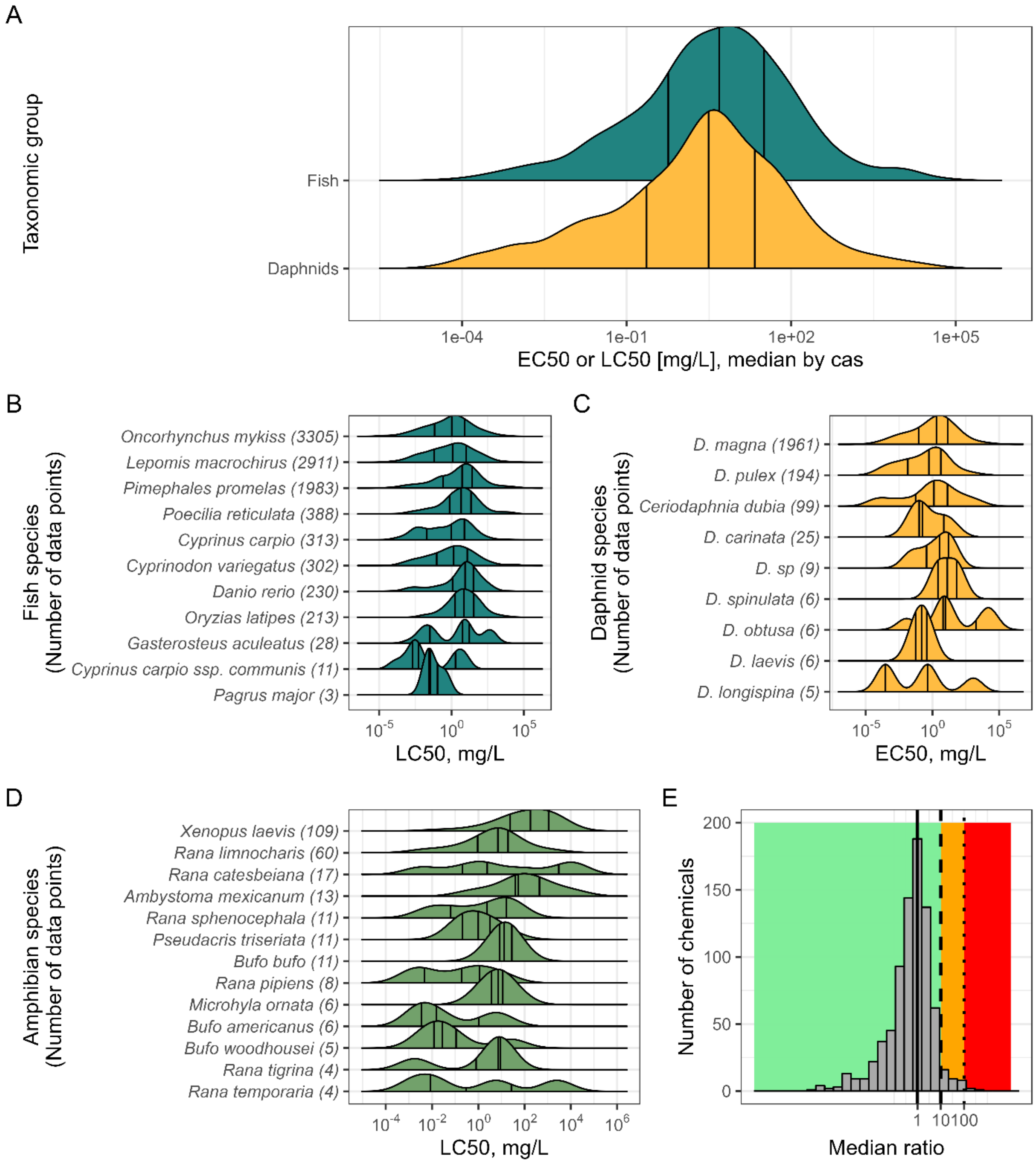


**Figure S2: (A) Distribution of median effect values ($EC_{50}$ and $LC_{50}$, summarized by chemical) for the two taxonomic groups, fish and daphnids.** Vertical lines in the distributions indicate quartile ranges. **Overview of the distributions of median $LC_{50}$ and $EC_{50}$ for fish (B), daphnids (C), and amphibians (D) separated by species (at least two data points, for fish and daphnids only non-summarized data points from chemicals in the final analysis). (E) Histogram of the distribution of ratios of median effect values for daphnids to fish.** A ratio of 1 indicates that for a specific chemical daphnids and fish are equally sensitive, while higher values correspond to a relatively higher sensitivity of fish (orange and red colors indicate ranges where fish are more sensitive than daphnids by a factor of at least 10 and 100, respectively), while values smaller than 1 correspond to higher sensitivity of daphnids (green background area). Figures B-D are limited to species with at least two data points. In panel C: *D.* = *Daphnia*.

**Correlation between number of data points per chemical and variability**

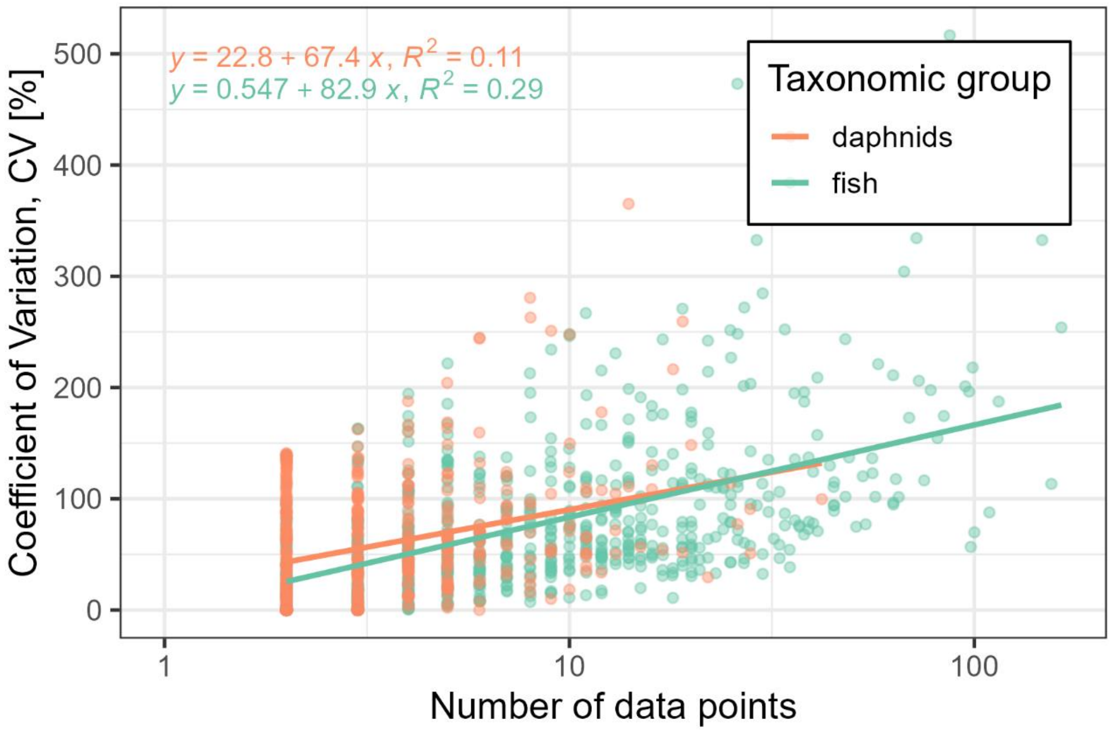


**Figure S3: Correlation of number of underlying data points per chemical and the coefficient of variation (CV) for the two taxonomic groups daphnids (orange) and fish (green).** The colored lines represent linear regression models represented by the equations above. Note that the CV of a single data point is 0, and thus not pictured.

### Correlation between fish and daphnid acute toxicity data

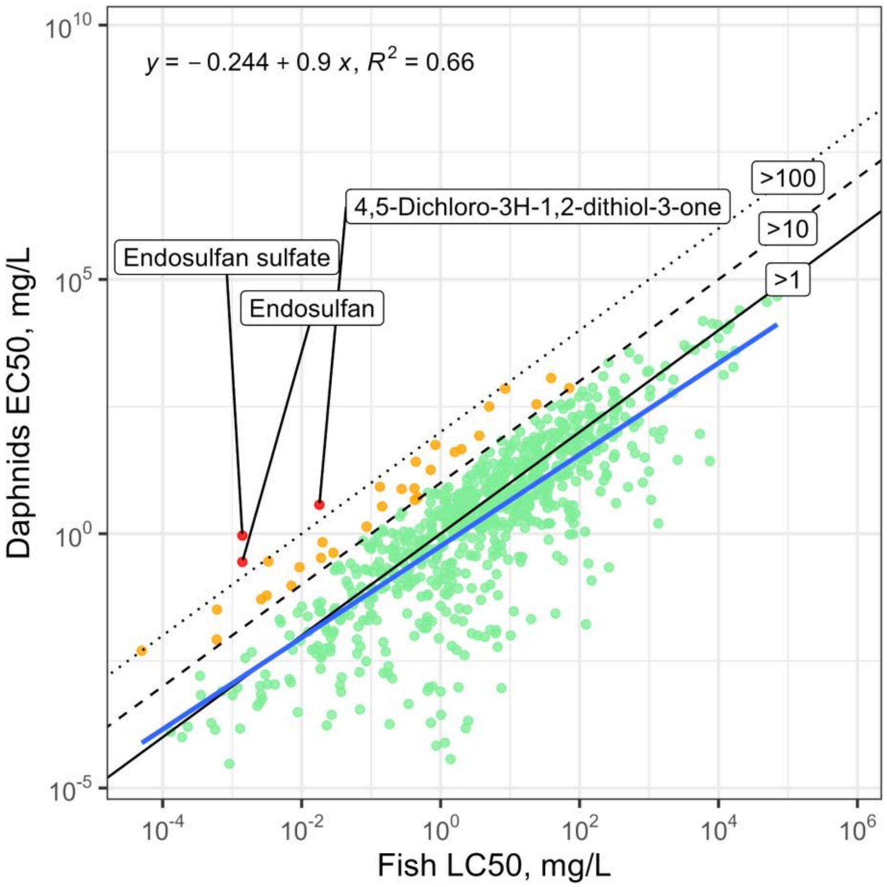


**Figure S4: Linear correlation between fish $LC_{50}$ (96 h) and daphnid $EC_{50}$ (48 h).** The black lines indicate ratio-limits of above 1, above 10 (orange dots) and above 100 (red dots). Only 3 compounds (irrespective of assigned MoA) have $R_{df}$ daphnid:fish ratios above 100: Endosulfan sulfate, endosulfan, and 4,5-dichloro-3H-1,2-dithiol-3-one.

### Significance of fish species variability

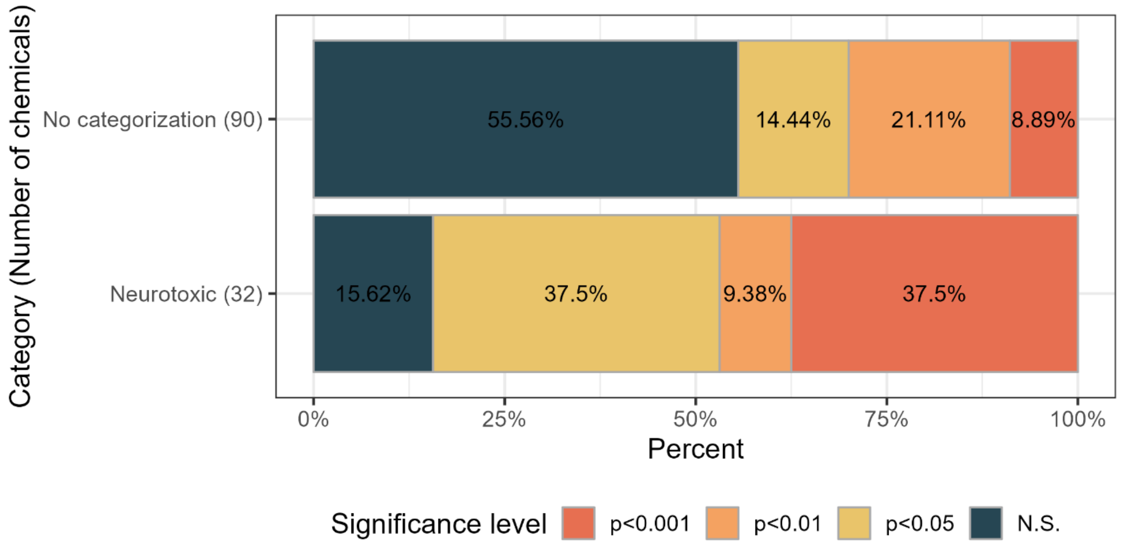


**Figure S5: Distribution of significance levels (Kruskal-Wallis rank sum test) for the inter-species differences for fish per chemical across the 90 chemicals with no categorization and 32 chemicals with a neurotoxicity label according to IRAC and with data for at least 4 tested species. (N.S. = not significant, p-value > 0.05).**

### Species-variability for neurotoxic and uncategorized compounds in daphnids

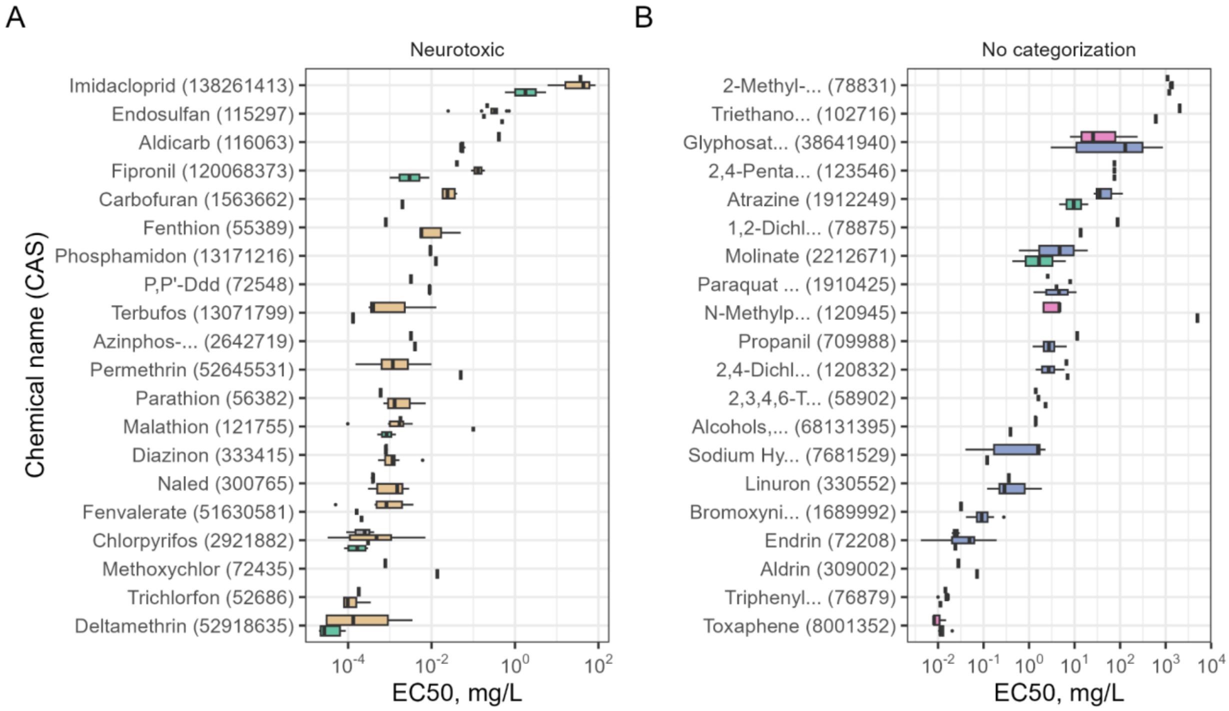


**Figure S6: Variability of $EC_{50}$ values across more than two species in daphnids for 20 randomly selected chemicals from the chemicals assigned (A) a neurotoxic MoA and (B) no categorization through IRAC.**

**Distribution of CVs for acute fish $LC_{50}$ values according to the analysis in Braunbeck et al. (2020): "*Development of an OECD Guidance Document for the Application of OECD Test Guideline 236 (Acute Fish Embryo Toxicity Test)*".**

In an earlier analysis, Braunbeck *et al.* (2020) applied more stringent filters to the fish acute toxicity data. They eliminated all fish $LC_{50}$ values that were higher than the modeled water solubility and where no data on water solubility was available, or when the log $P_{ow}$ value was above four and water concentration was not analytically confirmed, or when no data on log $P_{ow}$ were available. With this approach, the CVs range up to 150% with a median of approximately 20%. This estimate is lower, but also less robust, since it can be based only on 58 chemicals and it contains only two chemicals with a neurotoxic MoA according to the classification scheme applied in the present publication.

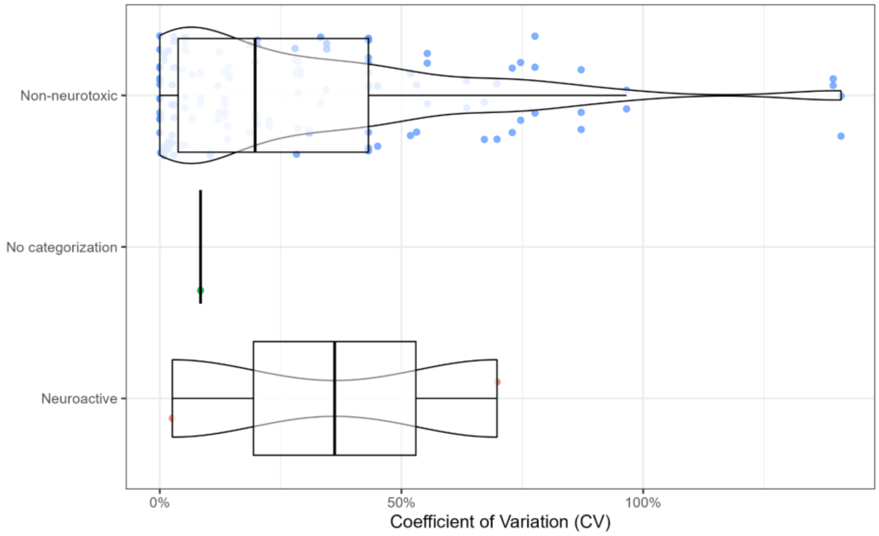


**Figure S7: Distribution of CVs for chemicals fish $LC_{50}$ values, filtered according to the more stringent criteria described in by Braunbeck et al. 2020** (compared to the approach applied this analysis (see Figure 4 in the main manuscript).

**Table S3: List of all chemicals in the analysis with $R_{df}$ values above 10. CV = Coefficient of variation. Categories for TEST, ASTER, IRAC, EFSA, and KRAMER have at times been simplified from the original sources. Chemical names with an asterisk (*) were not given in the dataset and have been added manually from the ECHA registration dossier website according to the CAS number.**

| Chemical Name | CAS Number | $R_{df}$ Daphnid /fish median ratio | CV fish [N] | CV daphnids [N] | TEST | ASTER | IRAC | EFSA | KRAMER |
|---|---|---|---|---|---|---|---|---|---|
| Endosulfan sulfate | 1031078 | 657,1428571 | [1] | [1] | Non-Neurotoxic | Neurotoxicant: Cyclodiene-Type | No Categorization | No Categorization | Non-Neurotoxic |
| Resmethrin | 10453868 | 19,67557252 | 74,87126937 [51] | 112,1590907 [4] | Neurotoxicity | Neurotoxicant: Pyrethroid | Sodium Channel Modulators | No Categorization | Neuroactive |
| Maleic acid | 110167 | 63,22 | [1] | 0,044739436 [2] | Non-Neurotoxic | Non-Neurotoxic | No Categorization | No Categorization | Non-Neurotoxic |
| Fenoxaprop-P | 113158400 | 27,49454545 | [1] | 131,6578398 [2] | Non-Neurotoxic | Non-Neurotoxic | No Categorization | No Categorization | Non-Neurotoxic |
| Endosulfan | 115297 | 200 | 187,3995313 [115] | 51,55554666 [19] | Neurotoxicity | Neurotoxicant: Cyclodiene-Type | Gaba Gated Clc Blockers Cyclodiene Organochlorines | No Categorization | Neuroactive |
| 4,5-Dichloro-3H-1,2-dithiol-3-one | 1192525 | 205,8611111 | 221,67585 [5] | 141,0015384 [2] | Non-Neurotoxic | Non-Neurotoxic | No Categorization | No Categorization | No Categorization |
| Cyazofamid | 120116883 | 85,9939759 | [1] | 63,15664771 [2] | Non-Neurotoxic | Non-Neurotoxic | No Categorization | No Categorization | Non-Neurotoxic |
| Anthracene | 120127 | 13,33333333 | 113,7859415 [14] | 45,60452924 [3] | Non-Neurotoxic | Non-Neurotoxic | No Categorization | No Categorization | Non-Neurotoxic |
| Diallyl phthalate | 131179 | 59,09090909 | [1] | [1] | Non-Neurotoxic | Non-Neurotoxic | No Categorization | No Categorization | No Categorization |
| Captan | 133062 | 62,68656716 | 55,8994861 [21] | [1] | Non-Neurotoxic | Non-Neurotoxic | No Categorization | No Categorization | Non-Neurotoxic |
| Naphthenic acids, copper salts* | 1338029 | 10,81081081 | 101,7112136 [4] | [1] | No Categorization | No Categorization | No Categorization | No Categorization | No Categorization |
| Pyrithione zinc* | 13463417 | 19,42675159 | 20,7177783 [2] | 72,37682995 [3] | No Categorization | No Categorization | No Categorization | No Categorization | No Categorization |
| Dipropyl 2,5-pyridinedicarboxylate | 136458 | 25 | 44,90502094 [4] | [1] | Non-Neurotoxic | Non-Neurotoxic | No Categorization | No Categorization | No Categorization |
| Antimycin A | 1397940 | 100 | 196,3437042 [97] | [1] | Non-Neurotoxic | Non-Neurotoxic | No Categorization | No Categorization | No Categorization |
| Bethoxazin | 163269305 | 14,80701754 | 76,91336918 [2] | [1] | Non-Neurotoxic | Non-Neurotoxic | No Categorization | No Categorization | No Categorization |
| Pyraclostrobin | 175013180 | 17,57894737 | 212,7959292 [8] | 84,04152969 [3] | AChE Inhibition | Non-Neurotoxic | No Categorization | No Categorization | Non-Neurotoxic |
| Propachlor | 1918167 | 18,57142857 | 110,0868456 [5] | 35,66341193 [3] | Non-Neurotoxic | Non-Neurotoxic | No Categorization | No Categorization | Non-Neurotoxic |

| Chemical Name | CAS Number | $R_{df}$ Daphnid /fish median ratio | CV fish [N] | CV daphnids [N] | TEST | ASTER | IRAC | EFSA | KRAMER |
|---|---|---|---|---|---|---|---|---|---|
| 2-(Dimethylamino)ethyl acrylate | 2439352 | 83,64705882 | [1] | [1] | Non-Neurotoxic | Non-Neurotoxic | No Categorization | No Categorization | No Categorization |
| 2,2'-(1-Methyltrimethylenedioxy)bis(4-methyl-1,3,2-dioxaborinane) | 2665136 | 10,28169014 | [1] | [1] | Non-Neurotoxic | Non-Neurotoxic | No Categorization | No Categorization | No Categorization |
| cis-Captafol | 2939802 | 23,88888889 | 195,1891022 [8] | [1] | Non-Neurotoxic | Non-Neurotoxic | No Categorization | No Categorization | No Categorization |
| Fluchloralin | 33245395 | 34 | 74,20541225 [19] | 24,95670992 [2] | Non-Neurotoxic | Non-Neurotoxic | No Categorization | No Categorization | Non-Neurotoxic |
| Carboxin | 5234684 | 23,44444444 | 70,60060882 [17] | [1] | Non-Neurotoxic | Non-Neurotoxic | No Categorization | No Categorization | Non-Neurotoxic |
| Trinitroglycerin | 55630 | 23,05764411 | 46,94950048 [22] | [1] | Non-Neurotoxic | Non-Neurotoxic | No Categorization | No Categorization | No Categorization |
| Lindane | 58899 | 15,98837209 | 516,5965756 [87] | 108,6446596 [16] | Non-Neurotoxic | Non-Neurotoxic | No Categorization | Neurotoxic | Non-Neurotoxic |
| Dieldrin | 60571 | 23,65591398 | 206,0596056 [73] | 28,79288202 [8] | Neurotoxicity | Neurotoxicant: Cyclodiene-Type | No Categorization | Neurotoxic | Neuroactive |
| Salicylic acid | 69727 | 29,48717949 | [1] | 42,35334923 [3] | Non-Neurotoxic | Non-Neurotoxic | No Categorization | No Categorization | Non-Neurotoxic |
| Flucythrinate | 70124775 | 13,83333333 | 72,05875844 [15] | [1] | Non-Neurotoxic | Neurotoxicant: Pyrethroid | Sodium Channel Modulators | No Categorization | Neuroactive |
| Endrin | 72208 | 53,30578512 | 368,6386557 [80] | 107,6630699 [12] | Neurotoxicity | Neurotoxicant: Cyclodiene-Type | No Categorization | No Categorization | Neuroactive |
| Sethoxydim | 74051802 | 25,21875 | 48,14913395 [7] | 132,3087038 [2] | Non-Neurotoxic | Non-Neurotoxic | No Categorization | No Categorization | Non-Neurotoxic |
| 2-(4,5-Dihydro-4-methyl-4-(1-methylethyl)-5-oxo-1H-imidazol-2-yl)-3-pyridinecarboxylic acid with 2-propanamine (1:1) | 81510830 | 14,58333333 | 88,78285523 [7] | 100,8654269 [3] | Non-Neurotoxic | Non-Neurotoxic | No Categorization | No Categorization | No Categorization |
| Allantoin* | 82027596 | 66,30952381 | 128,3592011 [9] | [1] | No Categorization | No Categorization | No Categorization | No Categorization | No Categorization |
| Dithiopyr | 97886458 | 11,06382979 | 94,28402169 [5] | [1] | Non-Neurotoxic | Non-Neurotoxic | No Categorization | No Categorization | Non-Neurotoxic |

**Protectiveness of daphnid data for amphibian toxicity**

Using the EnviroTox DB for an analysis of the sensitivity of the acute daphnid $EC_{50}$ values relative to the available acute amphibian $LC_{50}$ and $EC_{50}$ values provides similar results to the daphnids-to-fish comparison: For the majority of compounds (based on 93 chemicals, irrespective of neurotoxicity and MoA), daphnids are protective for amphibian acute toxicity. Since no OECD TG is available for acute amphibian toxicity and since data availability is more limited for amphibians, acute $LC_{50}$ and $EC_{50}$ values for mortality, mobility, and development at 48 and 96 hours were extracted where the effect value did not exceed five times the estimated water solubility and the effect type did not include the terms “egg”, “embryo”, “larva” or “fetus. This data was compared with the daphnids data, which were filtered in the same way as for the daphnids to fish ratio analysis.

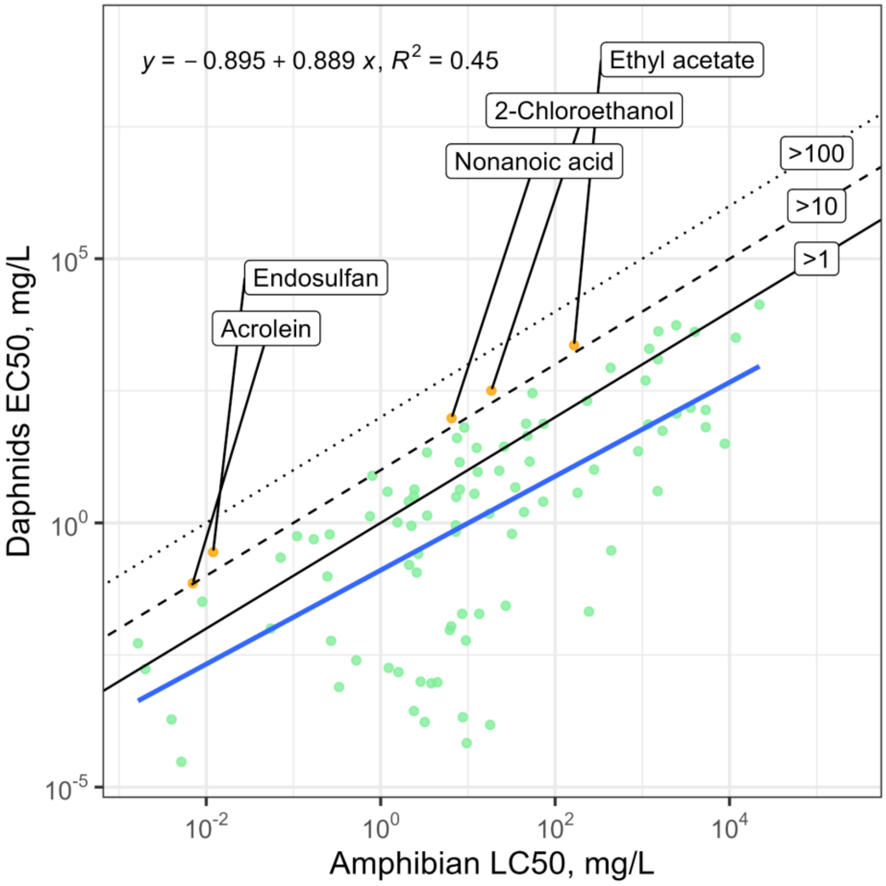


**Figure S8: Correlation analysis of daphnids acute toxicity data and amphibian acute toxicity data.** Colored points represent a chemical where both daphnid and amphibian data exists (N=93), color indicates the median ratio between these two groups (green: ≤ 10, yellow: > 10). Compounds with median ratios above 10 are labeled with the chemical name.

**Comparison of different data matching approaches**

There are different ways the daphnid and fish data can be summarized and matched with each other. Above, the $EC_{50}/LC_{50}$ values by CAS number were summarised, taking the median across all species, which results in one datapoint for each chemical that is represented in both the daphnid and the fish dataset ("By CAS"). This approach may be justified considering that the current OECD TG allows the use of multiple species and there is no robust knowledge on species-specific sensitivity to the different chemicals on the market. Moreover, from the available data it is not clear to what extent the selection of different species contributes to the overall variability in $LC_{50}$ and $EC_{50}$ data. For example, it appears that for malathion and trichlorfon the species-specific $LC_{50}$ distributions are relatively distinct, while they strongly overlap for fenitrotion and carbofuran. Thus, from a regulatory perspective, knowledge of taxonomic sensitivity differences may be most important. Moreover, from a scientific point of view, the "by CAS" grouping approach used throughout this study is most robust regarding individual $LC_{50}$ or $EC_{50}$ outliers. Additionally, this avoids bias for chemicals for which repeated tests with several different fish and daphnid species are available that have a stronger influence on the median ratio than other chemicals, which have been tested in fewer species.

Alternatively, one can summarize the data by chemical and species, resulting in several data points where a chemical was tested on multiple species ("By CAS and species"). Accordingly, this approach is highly influenced by more extreme values and outliers (*i.e.,* in terms of very low or very high (seemingly or true) species-specific $LC_{50}$ or $EC_{50}$ values).

A third approach takes these data and bases the median per chemical on these already summarized effect values (median per species and chemical → median per chemical; "By CAS, species, and CAS"). This approach gives equal weight to all tested species compared to the "By CAS", where species on which more datapoints are available have a higher impact on the $R_{df}$.

The fourth, and most granular, method of data matching is to pair every individual data point that was recorded for a chemical in the fish and in the daphnid dataset against each other, leading to an exponential increase in ratios and a strong bias towards chemicals (and species) that have been tested extraordinarily often ("All-to-all").

Previously, we compared the four approaches described above and how the distribution of median ratios across brackets depended on them (Figure S9, Figure S10, and Table S4).

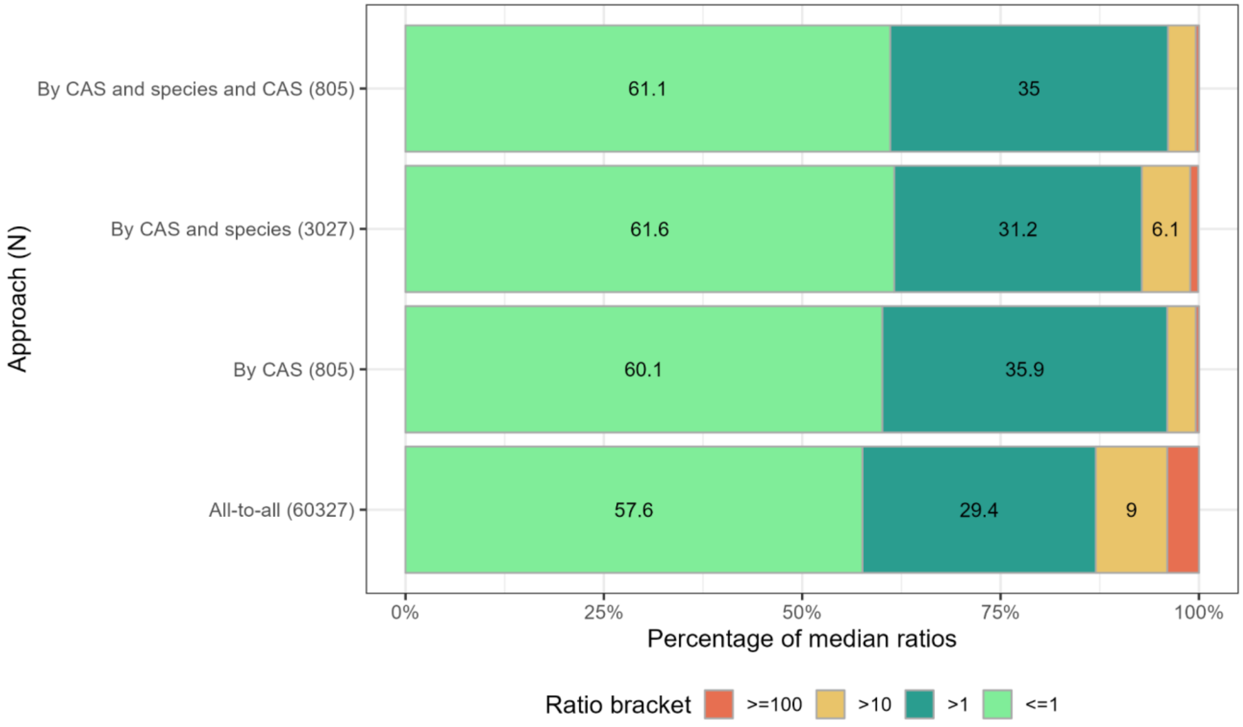


**Figure S9: Comparison of median ratios resulting from different approaches of summarizing and matching the fish and the daphnid data.**

Overall, the outcome is in line with the intuition that all-to-all matching and summarizing by CAS and species results in a higher percentage of ratios above 10 and above 100, respectively. This is to be expected since these approaches are more susceptible to outliers and extreme values, and since it is not possible to validate every individual data point in the dataset, the more conservative approach of analyzing the data summarized and matched by CAS was taken. These observations are mirrored in the distribution of median ratios visualized in the ridgeline plot of Figure S9. The all-to-all median ratios are distributed more broadly, as is expected as a result of the larger number of data points and matches resulting in more extreme values, while the distributions for the other approaches appear fairly similar. Nevertheless, the percentage of ratios above 10 appears fairly similar between these approaches, ranging from 3.5% to 9% (Table S4 below), which reassures that the overall conclusion from our analysis based on “By CAS” grouping is suitably robust.

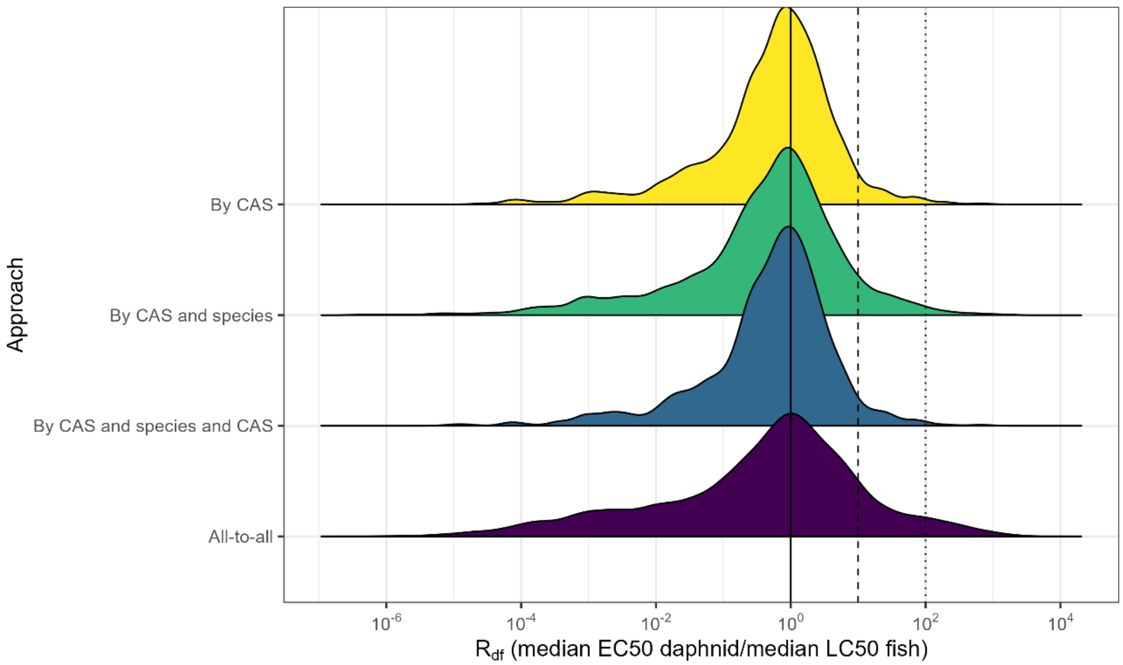


**Figure S10: Ridgeline plot of the distribution of median ratios resulting from different approaches to summarizing and matching the fish and daphnid data.**

**Table S4: Comparison of median ratios through different approaches of summarizing and matching the fish and daphnid data. Description of the different approaches are in the text above.**

| | | **Median ratio brackets (% and total N)** | | | |
|---|---|---|---|---|---|
| **Approach** | **Total N** | **% ≤1 (N)** | **% >1 (N)** | **% >10 (N)** | **% ≥100 (N)** |
| **All-to-all** | 60327 | 57.6 (34777) | 29.4 (17718) | 9 (5430) | 4 (2402) |
| **By CAS** | 805 | 60.1 (484) | 35.9 (289) | 3.6 (29) | 0.4 (3) |
| **By CAS and species** | 3027 | 61.6 (1866) | 31.2 (945) | 6.1 (185) | 1 (31) |
| **By CAS and species and CAS** | 805 | 61.1 (492) | 35 (282) | 3.5 (28) | 0.4 (3) |
